# Placental Iron Utilisation in Fetal Growth Restriction: Alterations in Mitochondrial Heme Synthesis and Iron-Sulfur Cluster Assembly Pathways

**DOI:** 10.1101/2025.06.09.658195

**Authors:** Veronica B Botha, Heather C Murray, Siddharth Acharya, Kirsty G Pringle, Roger Smith, Joshua J Fisher

**Affiliations:** School of Biomedical Sciences and Pharmacy, University of Newcastle, Callaghan, New South Wales, Australia; School of Medicine and Public Health, University of Newcastle, Callaghan, New South Wales, Australia; Precision Medicine Program, Hunter Medical Research Institute, New Lambton Heights, New South Wales, Australia; Women’s Health Research Program, Hunter Medical Research Institute, New Lambton Heights, New South Wales, Australia; Mothers and Babies Research Program, Hunter Medical Research Institute, New Lambton Heights, New South Wales, Australia

## Abstract

Fetal growth restriction (FGR) affects ∼10% of pregnancies worldwide and is often associated with placental insufficiency. Iron is essential for maternal haematopoietic adaptations and placental processes such as mitochondrial iron–sulfur (Fe-S) cluster assembly, heme synthesis, and erythropoiesis. This study aimed to characterise iron transport and downstream utilisation in FGR. Placental tissues from term uncomplicated (n=19) and FGR (n=18) pregnancies were analysed. Maternal iron status was retrospectively assessed from clinical records. Placental mRNA and protein expression of iron-dependent pathways were analysed via RT-qPCR, LC-MS, and western blotting. Placental iron content was assessed histologically, and heme levels were measured by activity assay. FGR pregnancies showed significantly elevated maternal serum ferritin and lower red cell distribution width, although remained within normal clinical values. Placental iron uptake transporters TFRC and DMT1 were significantly upregulated, while the iron exporter to the fetus, ferroportin, was reduced, indicating increased iron retention in the FGR placenta. Despite altered transporter expression, Fe^3^⁺ iron levels were unchanged, suggesting iron utilisation over storage. Subsequent investigations identified reduced mitochondrial Fe-S synthesis components (FDXR, FDX2, NDUFAB1, HSPA9), and a prioritisation of mitochondrial and cytosolic heme synthesis enzymes in FGR. Protein levels of haemoglobin subunits (HBG1, HBG2, HBB, HBA1) and erythrocyte membrane markers (EPB41, EPB42, SPTA1, SPTB, ANK1) were decreased. These findings reveal a compensatory response in FGR placentae, with increased iron uptake and utilisation favouring heme synthesis over Fe-S cluster formation, possibly to support oxygen handling under poor placental vascularisation and reduced fetal oxygenation, with potential consequences for mitochondrial energy metabolism.

**Key Points:**

- Iron plays a critical role in placental function, while iron-dependent pathway components are well-characterised, their integrated response and adaptive reprogramming in fetal growth restriction (FGR) remain poorly understood.
- In FGR, maternal iron status was unchanged, however, placental iron uptake proteins were increased and ferroportin reduced, suggesting that the placenta retains iron.
- FGR placentae showed altered de novo mitochondrial iron-sulfur cluster (Fe-S) formation and a bottleneck in late-stage Fe-S cluster assembly.
- This shift in Fe-S synthesis prioritises mitochondrial and cytosolic heme synthesis pathways, consistent with increased heme utilisation and breakdown.
- Globin subunits were lower in protein abundance and impaired placental erythrocyte structure in FGR.
- Dysregulation of erythrocyte membrane proteins in FGR placentae suggests altered erythrocyte structure, potentially representing an adaptive response to inadequate vascularisation, attempting to optimise oxygen delivery to the fetus.

## Introduction

Fetal growth restriction (FGR) occurs in 10% of pregnancies globally and is associated with increased perinatal and neonatal morbidity and mortality (Miller *et al*., 2008). FGR is defined by the inability of the fetus to achieve its genetically predetermined growth potential relative to its gestational age (Gordijn *et al*., 2016). Although maternal and fetal complications are recognised causes of FGR, 60-70% of FGR cases are associated with placental insufficiency, where the placenta fails to adequately deliver nutrients and oxygen, arising from inadequate uterine spiral artery remodelling (Brooker *et al*., 2025; Higashijima *et al*., 2013; Tang *et al*., 2017). In FGR placentae, the cellular landscape is altered, with impaired cytotrophoblast maturation and decreases in trophoblast volume. This results in a reduction in the number of placental villi and a decreased placental diameter (Sun *et al*., 2020). Consequently, the surface area available for oxygen and nutrient exchange is diminished, limiting substrate transfer to the fetus (Zygmunt *et al*., 2003). In response, the fetus adapts by redistributing blood flow, prioritising essential fetal organs, such as the brain and heart, resulting in asymmetrical growth and slowing the fetal growth rate (Miller *et al*., 2008; Roberts *et al*., 2020; Wu *et al*., 2023). The placenta is a highly vascularised, potent haematopoietic organ. The placenta performs primitive haematopoiesis during embryogenesis, establishing a large reservoir of haematopoietic stem cells (HSCs) that not only support erythrocyte activity during early gestation, but also provide progenitor cells for the fetal liver and bone marrow when haematopoiesis transitions away from the placenta (Gekas *et al*., 2010; Palis & Segel, 1998). Hematopoietic function is critically reliant on iron availability, as iron is essential for erythropoiesis and mitochondrial functions that facilitate the biosynthesis of iron-sulfur clusters (Fe-S) and subsequent heme synthesis in erythroid cells (Chiabrando *et al*., 2014; Hentze *et al*., 2004; Mastrogiannaki *et al*., 2009).

Iron is an essential micronutrient for both fetoplacental development and mitochondrial bioenergetics (Dev & Babitt, 2017; Guo *et al*., 2019). During pregnancy, the requirement for iron alters as gestation advances. In early pregnancy, placental development occurs in a physiologically hypoxic environment, with relatively low iron demands (Guo *et al*., 2016). This low-oxygen environment stabilises hypoxia inducible factors (HIFs), which transcriptionally activate genes involved in oxygen and iron regulation, such as erythropoietin (EPO), hepcidin and erythroferrone (Semenza, 2014). In the first trimester, rising EPO levels support both maternal erythropoiesis and transient placental hematopoietic activity (Cindrova-Davies & Sferruzzi-Perri, 2022; Delaney *et al*., 2021; Fairchild Benyo & Conrad, 1999). Consequently, hepcidin is suppressed, enhancing maternal iron mobilisation and supporting the increase in maternal iron demands to meet increased cardiovascular, renal, and haematological adaptations, as well as fetoplacental development (Zaugg *et al*., 2022). These adaptations require sufficient iron for enhanced haematopoiesis and haemoglobin synthesis. Subsequently, this supports optimal oxygen transfer, thereby sustaining fetal development, while also establishing fetal iron reserves in the later stages of gestation (Cao & Fleming, 2016; Roberts et al., 2020). Maternal iron deficiency, anaemia and excess iron alike have been implicated in adverse pregnancy outcomes, including FGR (Andrews, 2000), highlighting the important role of iron in healthy pregnancies.

Iron transport during pregnancy occurs unidirectionally from the mother to the fetus, primarily via transferrin-bound iron in the maternal circulation (McArdle *et al*., 2003). At the maternal placental interface, transferrin receptor 1 (TfR1/TFRC) binds the maternal transferrin-iron complex, inducing endocytosis (McArdle *et al*., 2011; Roberts *et al*., 2020). Within the endosome, the acidic pH induces a conformational change in TfR1, releasing ferric (Fe^3+^) iron. Following the conversion of Fe^3+^ to ferrous (Fe^2+^) iron, iron is transported into the cytosol via divalent metal transporter 1 (DMT1) (McArdle *et al*., 2011; Roberts *et al*., 2020; Waldvogel-Abramowski *et al*., 2014). Once in the cytosol, iron has three potential fates, it is either: (i) stored in ferritin, (ii) shuttled to the mitochondria for iron-sulfur (Fe-S) cluster biogenesis and heme synthesis, or (iii) transported across the basolateral (fetal) membrane via ferroportin for entry into the fetal circulation following reoxidation to Fe^3+^ (McArdle *et al*., 2011; Roberts *et al*., 2020). Approximately 20-50% of cellular iron is directed towards the mitochondria to support two critical pathways, the biosynthesis of Fe-S clusters and heme (Qi *et al*., 2023; Roberts *et al*., 2020)

Fe-S clusters are diverse molecular structures that perform various biological functions within the cell. Specifically, Fe-S clusters play a role within the mitochondrial electron transport chain (ETC), via redox reactions that transfer electrons and generate a proton gradient essential for ATP synthesis (Mostajabi Sarhangi & Matyushov, 2023; Paul *et al*., 2017). De novo Fe-S cluster synthesis begins with the formation of [2Fe-2S] clusters, which serve as essential precursors in the Fe-S cluster pathway. These de novo [2Fe-2S] clusters undergo further processing to generate more complex [3Fe-4S] and [4Fe-4S] clusters downstream. These late-stage [3Fe-4S] and [4Fe-4S] clusters are critical for the function and stability of the mitochondrial ETC, particularly in complexes I, II and III, where they facilitate efficient electron transfer during oxidative phosphorylation (Maio & Rouault, 2020; Mostajabi Sarhangi & Matyushov, 2023; Read *et al*., 2021). Additionally, the terminal enzyme in heme biosynthesis, ferrochelatase (FECH), requires [2Fe-2S] clusters for its activity, linking Fe-S cluster synthesis with heme production. Heme is essential for oxygen transport through its incorporation into haemoglobin and cytochromes, which facilitate electron transfer in the ETC, particularly complexes II, III, and IV (Anderson & Frazer, 2017; Kim *et al*., 2012).

The primary utilisation of iron in fetal development is in erythropoiesis (Cerami, 2017), and heme stimulates the production of α- and β-like globin genes with distinct haemoglobin profiles expressed at specific stages of gestation (Kim *et al*., 2012; Moksnes *et al*., 2022). Embryonic (ζ2ε2) and fetal (α2γ2) haemoglobin exhibit higher oxygen affinity compared to adult (α2β2) haemoglobin, demonstrated by a leftward shift in their oxygen dissociation curves. These haemoglobin variants exhibit higher oxygen affinities during early development, enabling adaptation to lower oxygen availability and efficient maternal–fetal oxygen transfer (Manning *et al*., 2020).

Despite the well-described crucial role of iron in placental and fetal development, the specific alterations in iron-dependent pathways in FGR placentae remain uncharacterised. The complex interplay between iron transport, Fe-S cluster assembly, and heme synthesis is particularly relevant in the context of FGR, where placental insufficiency and impaired oxygen delivery are defining clinical characteristics. We have therefore examined genes and proteins involved in placental iron transport, as well as iron-related mitochondrial proteins involved in Fe-S assembly, heme synthesis, and placental erythropoiesis in FGR compared to healthy placentae. Collectively, these results provide molecular insights into how altered iron transport and iron-dependent pathways may contribute to mitochondrial dysfunction and placental insufficiency observed in FGR.

## Methodology

### Ethical Approval

Ethical approval for this research was obtained from the University of Newcastle Human Research Ethics Committee (H-382-0602), Hunter New England Health Human Research Ethics Committee (02/06/12/3.13), and Site-Specific Assessment (SSA/15/HNE/291), in compliance with the Declaration of Helsinki. Informed consent was obtained from all participants.

### Maternal and Placental Sample Collection

Placental villous cores were collected from 19 healthy, uncomplicated and 18 FGR (percentiles: <3^rd^ n=7, <5^th^ n=2 and <10^th^ n=9) pregnancies within 45 minutes of birth as previously described by Fisher *et al*. (2024) and washed in ice-cold PBS to limit maternal blood contamination. Tissues were either preserved in formalin at 4°C overnight and stored in 0.1 M PBS with 0.05% sodium azide for histological analysis, or snap-frozen and stored at −80°C for later gene and protein analysis. FGR was diagnosed with consensus criteria as published by Gordijn *et al*. (2016). All placentae collected for this study met the eligibility criteria. Healthy term pregnancies were from singleton deliveries between 37-40 weeks of gestation. FGR cases were from singleton pregnancies delivered by caesarean section at 34-40 weeks of gestation with an estimated fetal weight <10th centile and birthweight confirmed <10^th^ centile. Exclusion criteria applied to all samples with premature membrane rupture, hypertension, diabetes, placental abnormalities, chorioamnionitis, immune deficiencies, infectious diseases, smoking, non-steroidal anti-inflammatory drug use, illicit drug use, and metformin use were not eligible within our study. Maternal blood samples collected as part of routine care were analysed retrospectively to assess clinical iron parameters. Reference ranges were obtained using New South Wales Health Pathology (NSWHP) Hospital Laboratory Clinical Projects Laboratory Manual (Brighton, 2019) in conjunction with Pregnancy and Laboratory Studies: A Reference Table for Clinicians (Abbassi-Ghanavati *et al*., 2009).

### Histological analysis of iron deposition by Prussian Blue staining

To detect Fe^3+,^ samples were embedded in paraffin and sectioned at a thickness of 4 µm using a Leica RM2135 Microtome (Leica Biosystems, Germany). Fixed slides were dewaxed in a series of xylene and ethanol submersions and then washed in MilliQ water. A subset of collected placental villous tissue slides (Control n=7 and FGR n=7; : <3^rd^ n=3, <5^th^ n=2, <10^th^ n=2) were stained with 10% (weight/weight) Prussian Blue (potassium hexacyanoferrate (II) trihydrate, Sigma-Aldrich, AUS) for 5 minutes, then 5% (weight/weight) Prussian Blue stain in 10% HCl for 30 minutes, and counterstained with nuclear fast red for 5 minutes (Sigma Aldrich, AUS) as described by Zaugg *et al*. (2024). Slides were then washed and dehydrated following gradient ethanol exposure of 70% and 100% for 3 minutes, finishing with a 10-minute xylene exposure. Slides were mounted with a coverslip and left to dry overnight. Slides were then imaged on Aperio GT 450 DX (Leica Biosystems, Germany) at 40X magnification. The intensity of the Prussian blue staining was quantified using H-scores obtained from Halo Quantitative Pathology analysis software (Indica Labs; USA). All images were processed using identical parameters to ensure valid comparisons between healthy and FGR placental samples.

### Placental RNA Preparation and Gene Expression Studies

Placental tissue was crushed on dry ice with liquid nitrogen and subsequently, lysed and homogenised with 1 mL Trizol® Reagent (Life Technologies; Thermo Fisher Scientific, AUS) using the Precellys24 instrument (Bertin Technologies, France) for two cycles of 30 seconds at 5000 × g, incubated on ice for 30 minutes, and then repeated. Samples were centrifuged for 10 minutes at 10,000 × g using the Microfuge 20R (Beckman Coulter, AUS). The remainder of the protocol was performed in line with the manufacturer’s instructions outlined in the Direct-zol^TM^ RNA mini-prep kit (Zymo Research, California, USA). Following quantification on the Nanodrop One (Thermo Scientific, AUS), 1000 ng of cDNA was synthesised using SuperScript^TM^ IV First-Strand Synthesis System (Thermo Fisher Scientific, AUS). RT-qPCR was conducted using the PowerUpTM SYBR Green Master Mix (Applied Biosystems, Thermo Fisher, AUS) with KiCqstart primers (Sigma Aldrich, AUS; Table 1). RT-qPCR was performed using the QuantStudio6 Flex Real-Time PCR system (Thermo Fisher Scientific, MA, USA). Relative gene expression was determined by normalising to the geometric mean of three housekeeping genes (β-Actin, TBP, and YWHAZ), which were quantified using the 2^-ΔΔCT^ method to assess relative gene expression within placental tissue samples.

**Table 1:**
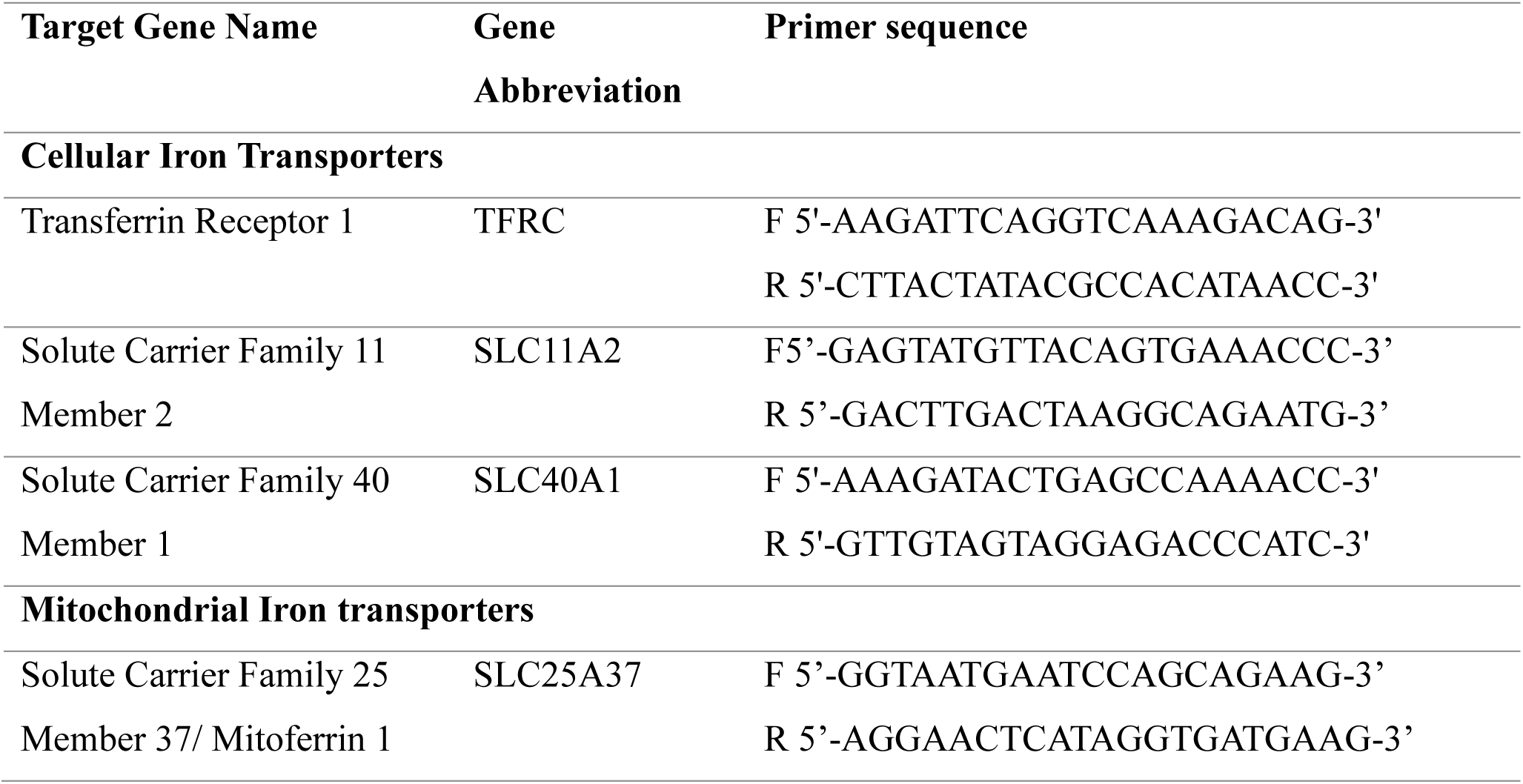
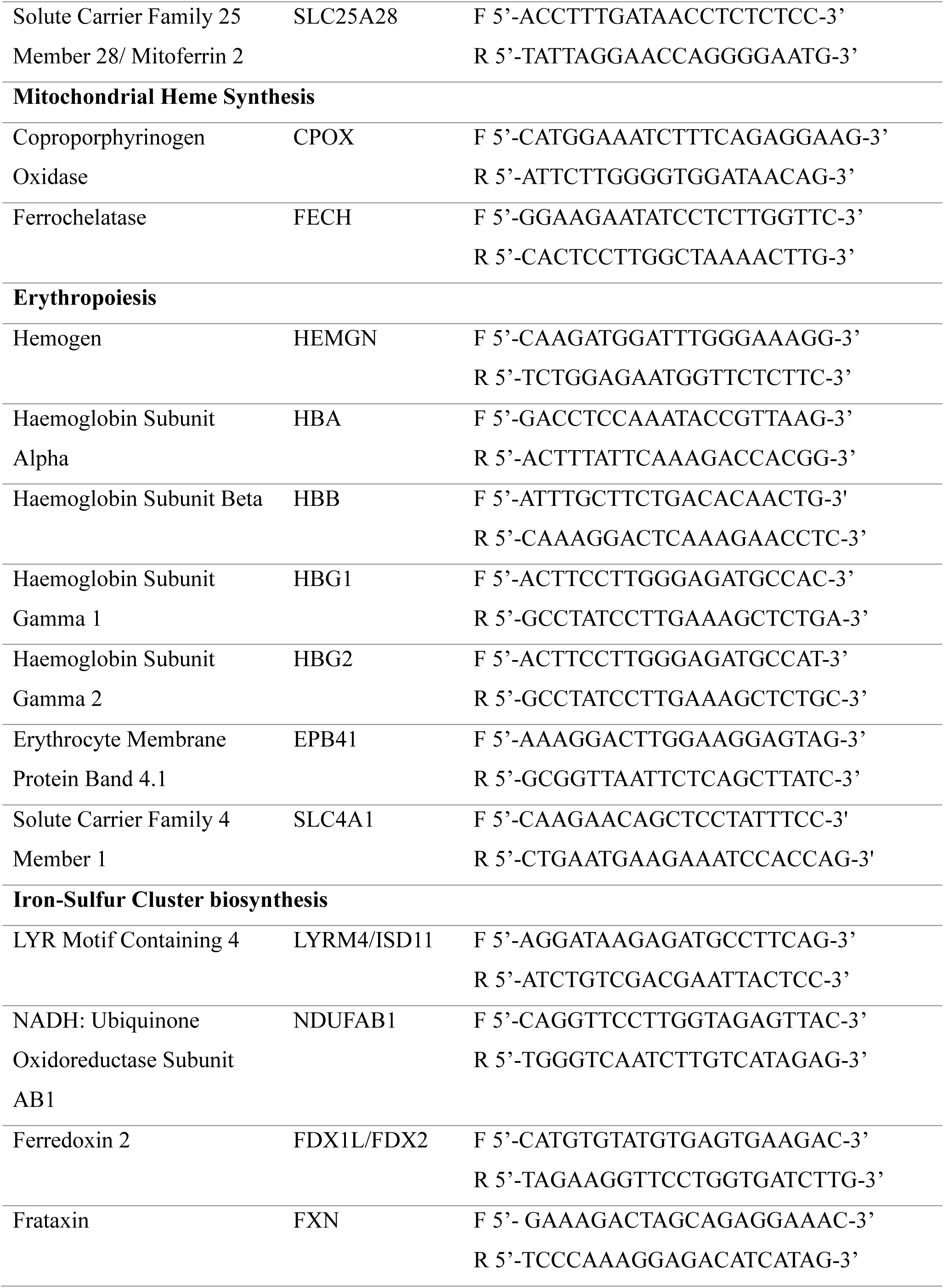
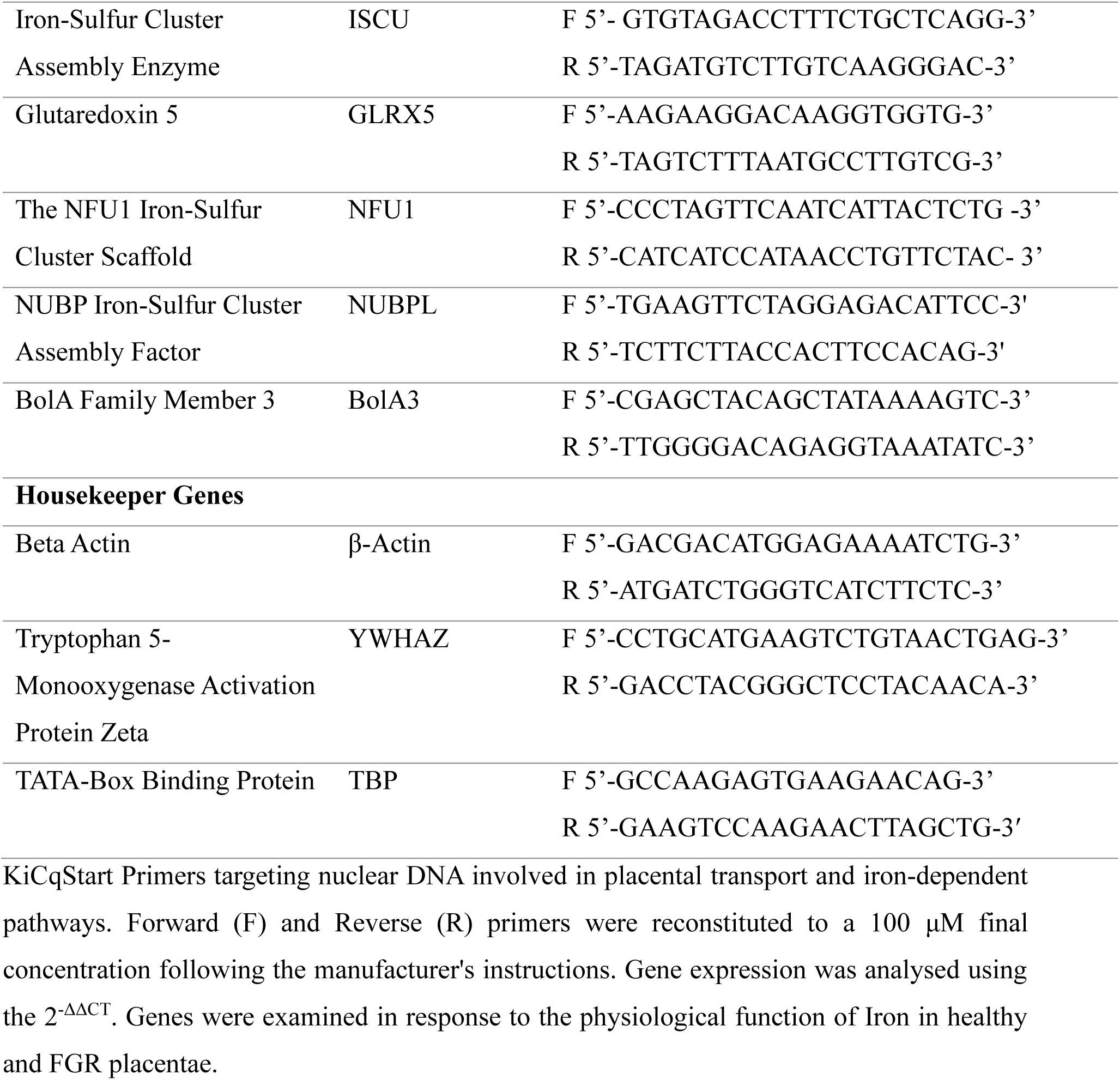
Kicqstart Primers for Gene Expression Studies of Targeted Iron Transporters and Intermediaries.

### Proteomics Studies

Placental tissue was crushed on dry ice and stored at −80°C until Liquid Chromatography-Mass Spectrometry (LC-MS) based proteomic analysis could be performed. LC-MS sample preparation followed previously published methodology by Mulhall *et al*. (2024), with placental tissue samples lysed in chilled 4% (weight/volume) sodium deoxycholate (SDC), 100 mM Tris HCl (pH.8.5) and passed through an 18G blunt-end needle before sonification. Samples were quantified and concentrations equalised to 200 μg/μl before digestion, alkylation and overnight trypsin incubation. Following trypsin exposure, peptides were diluted 1:1 with 100 mM Tris-HCl (ensuring SDC concentration < 1%) and vortexed briefly before adding 99% ethylacetate /1% Trifluoroacetic acid (TFA; 1:1) and thoroughly vortexed at 2000 rpm using ThermoMixer. Peptides were then desalted using styrene-divinylbenzene reverse-phase sulfonated StageTips.

Peptide separation was performed on an Aurora Ultimate C18 column (25 cm × 75μm, IonOpticks) using a 90-minute linear gradient from 3% to 41% of solvent B (acetonitrile in 0.1% formic acid) at a flow rate of 400 nL/min. MS analysis was conducted on an Eclipse mass spectrometer (Thermo Fisher Scientific, AUS), and MS scans were acquired over a mass range of 375-1400 m/z at a resolution of 60,000, with an automatic gain control (AGC) target of 100% and maximum injection time of 50 ms. MS/MS scans were acquired at a resolution of 15,000, with an AGC target of 100%, normalised collision energy of 30%, and dynamic maximum injection time. Raw data was analysed using Proteome Discoverer 2.5 (Thermo Fisher Scientific, AUS), outlined by Murray *et al*. (2023).

### Western Blotting

To validate our proteomics data, western blotting was performed. Protein was extracted from healthy (n=7) and FGR (n=7) placental tissue using 1 mL RIPA buffer (50 nM Tris-HCl (pH 7.4), 150 nM NaCl, 1% NP-P40, 1% sodium deoxycholate, 1% SDS, Complete Mini Protease Inhibitor Cocktail tablets (Roche Diagnostics, North Ryde AUS) and 1 nM PMFS). Samples were homogenised using Precellys24 and centrifuged as outlined previously. The supernatant was collected and stored at −80°C for subsequent western blotting analysis. Following Fisher *et al*. (2019) published protocol, samples (40 µg/µL) were loaded onto 12% Tris gels (Invitrogen, AUS), electrophoresed, and transferred to Polyvinylidene fluoride (PVDF) membranes. Membranes were blocked with Odyssey Intercept buffer (LICORBio, USA) for 1 hour, then incubated overnight with primary antibody, LYMR4 at 1:250 dilution (ab253001, Abcam, AUS) and Beta Actin (ab8226, Abcam, AUS) at a 1:1000 dilution. After washing with PBST, membranes were incubated for 1 hour with secondary antibodies anti-mouse IRDye®800CW and anti-rabbit IRDye®680LT (LICORBio, USA) at 1:20,000 dilution and washed with PBST. Membranes were imaged using the LI-COR Odyssey M system (LICORBio, USA) and analysed with Studio Image v5.2.

### Assessment of Heme

Placental heme concentrations were measured using a Heme Assay kit (MAK316, Sigma Aldrich, AUS). Following the manufacturing instructions, a total of 250 μL of water was added to a 96-well plate, creating a blank well. A standard well was created by mixing 50 μL of heme calibrator and 200 μL of water. Subsequently, 50 μL of homogenised term and FGR samples suspended in glycerol buffer (50 mM Tris HCl, 1 mM EDTA, 0.5% Glycerol) was mixed with 200 μL of reagent respectively. The plate was left to incubate for 5 mins at room temperature, and the absorbance of well contents was determined at 400 nm. All samples and standards were performed in duplicate, and total heme concentration was determined using the following equation:

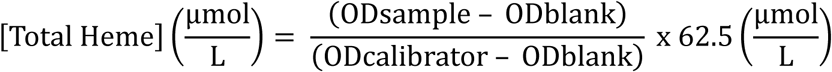

### Statistical Analysis

All data were analysed using GraphPad Prism 10.0. Outliers were identified and removed using Grubbs and ROUT tests, and data normality was assessed using the Shapiro-Wilk test. For normally distributed data, student t-tests were performed. For data that did not follow a normal distribution, Welch t-tests were conducted. Statistical significance was set at p < 0.05. John Hunter Hospital clinical data are presented with 95% confidence intervals (CI) and interquartile ranges (IQR). Gene expression data were presented as mean ± standard error of the mean (SEM), and histological H-scores, accounting for blue-stained iron deposits within placental tissue, were quantified using Halo Quantitative Pathology Analysis Software (Indica Labs, USA) and calculated using the following formula (Wen *et al*., 2024):

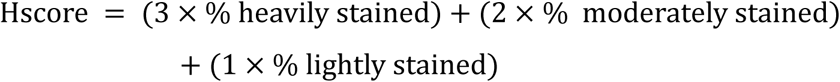

This formula weights the staining intensity, generating a composite score ranging from 0-300, reflecting both the extent and intensity of iron deposits. Proteomic analysis was expressed as log_2_-fold change (FC) to determine differential protein expression between healthy controls and FGR, visualised using heatmaps. The log_2_-fold was derived from the ratio between healthy term placentae to assess changes in FGR placentae, so a positive log_2_-FC indicates higher protein abundance in healthy control placentae, whereas a negative log_2_-FC indicates higher protein abundance in FGR placentae. Peptides with fewer than 2 Peptide Spectrum Matches (PSMs) were excluded from the analysis.

## Results

### Clinical Insights into Maternal Iron Bioavailability, Birth and Gestational Characteristics

To better understand the potential role of iron throughout gestation, we evaluated clinical characteristics encompassing maternal physical attributes, birth-related factors, and gestational characteristics (Table 2). Analysis included maternal parameters (age and BMI), birth outcomes (gestational age at delivery, birth weight, birth centile, placental weight, fetal sex, mode of delivery) and fetal biometry measurements following sonography assessment (estimated fetal weight (EFW%), abdominal circumference, femur length, head circumference, and amniotic fluid volume). FGR babies were significantly smaller (p<0.0001), with a median birth weight of 2415g (95% CI: 1760g-2356g), compared to healthy term babies, who had a median birth weight of 3555g (95% CI: 3346g-3676g). Additionally, FGR babies had smaller placentae (p<0.0001), with a median placental weight of 556g (95% CI: 395g-519g), compared to healthy term babies, who had a median placental weight of 689g (95% CI: 628g-757g). Furthermore, fetal biometric measurements, including abdominal circumference (p<0.0001), femur length (p=0.002), and head circumference (p=0.0007), were all significantly lower in FGR pregnancies. Although the difference in amniotic fluid volume was not statistically significant, FGR pregnancies had lower amniotic fluid volume (p=0.054) compared to healthy term controls.

**Table 2:**
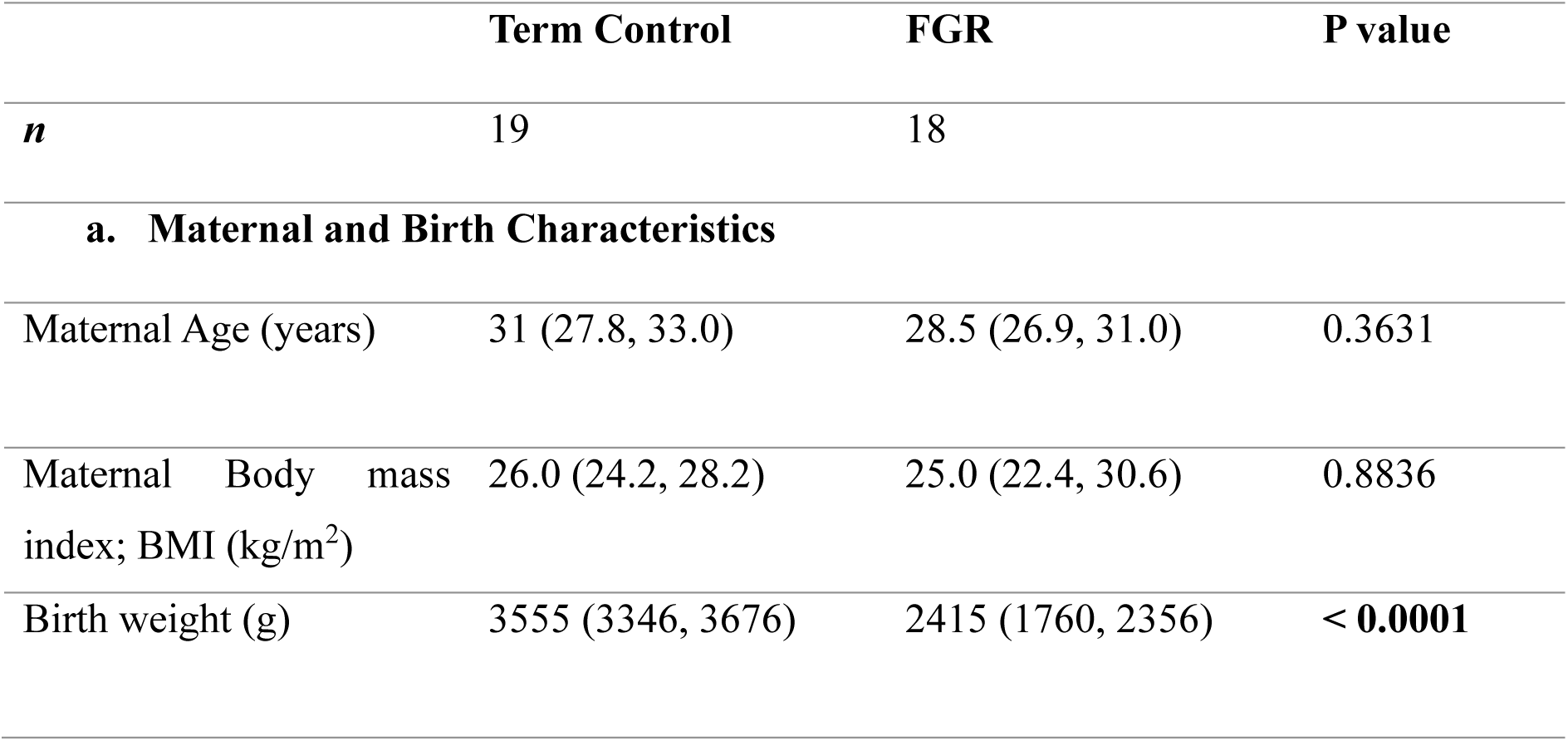
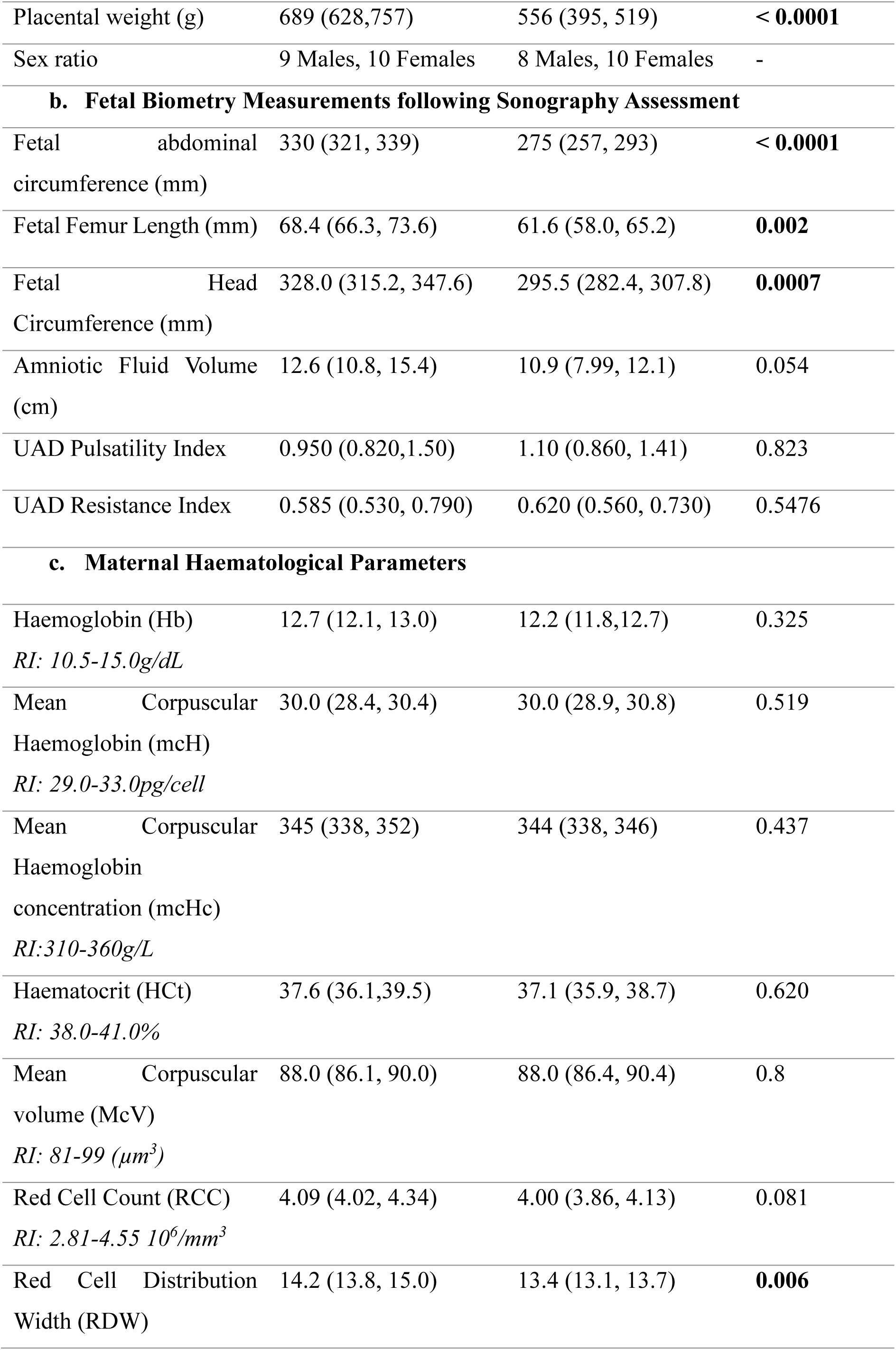
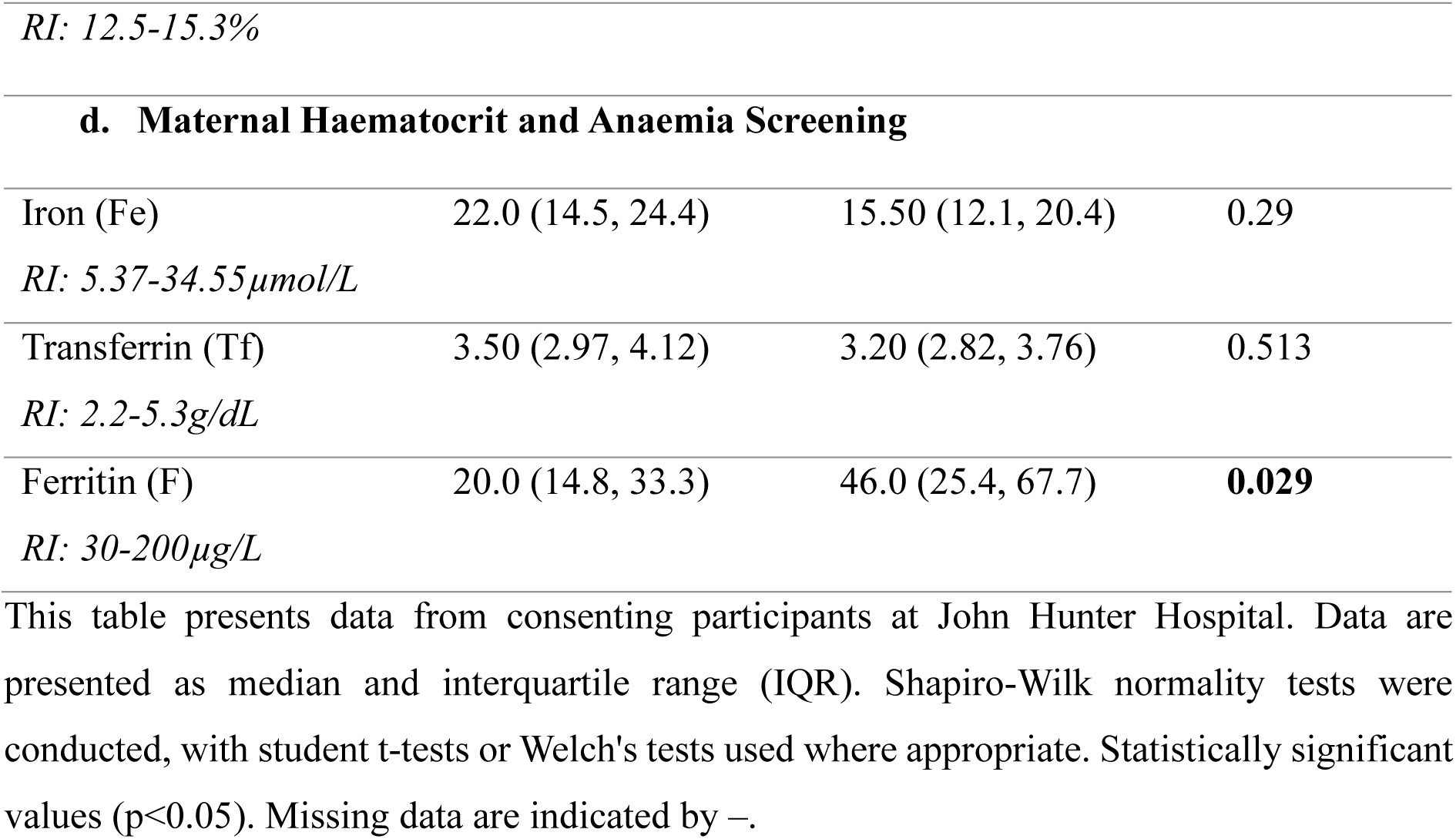
Maternal, Fetal and Birth Clinical Characteristics from Pregnancies Diagnosed as FGR And Healthy Term Controls.

Maternal haematological indices and maternal haematocrit and anaemia screening (Table 2) were performed on healthy and FGR pregnancies. No clinical or statistical significance was observed between FGR and healthy control groups in haemoglobin (Hb), mean corpuscular haemoglobin (MCH), mean corpuscular haemoglobin concentration (MCHC), mean corpuscular volume (MCV), red cell count (RCC), hematocrit (HCt), systemic iron levels and transferrin (Tf). Maternal ferritin levels were higher (p=0.029) and red cell distribution width (RDW) lower (p=0.006) in FGR cases compared to healthy control, however, both parameters fell within clinical reference ranges.

### Placental Iron Transport

Analysis of iron transporters between FGR and control placentae revealed significant alterations in gene expression and protein levels of iron import and export components. Transferrin Receptor 1 (TFRC) showed significantly increased mRNA expression (p<0.005, Figure 1A) and upregulated protein levels (−0.117 log_2_-FC; Figure 1D) in FGR placentae compared to healthy term placentae. Divalent Metal Transporter 1 (DMT1/SLC11A2) exhibited significantly increased mRNA expression in FGR (p<0.05, Figure 1B) compared with controls. In contrast, Ferroportin (FPN/SLC40A1; Figure 1C), the iron receptor responsible for transporting iron out of the placenta into fetal circulation, showed no significant difference in mRNA expression between FGR and healthy controls. FPN protein analysis (Figure 1D) revealed lower protein levels (0.646 log_2_-FC) in FGR placental villous tissue. These findings suggest FGR placentae are retaining iron for placental function. Following the clinical characterisation of maternal iron parameters and mRNA expression of iron transport into the placenta, the placental iron content between healthy and FGR placentae was assessed. Histological analysis by Prussian blue staining revealed intense blue regions with no statistical difference between healthy control (Figure 1F-H) and FGR (Figure 1I-K) placentae.

**Figure 1:**
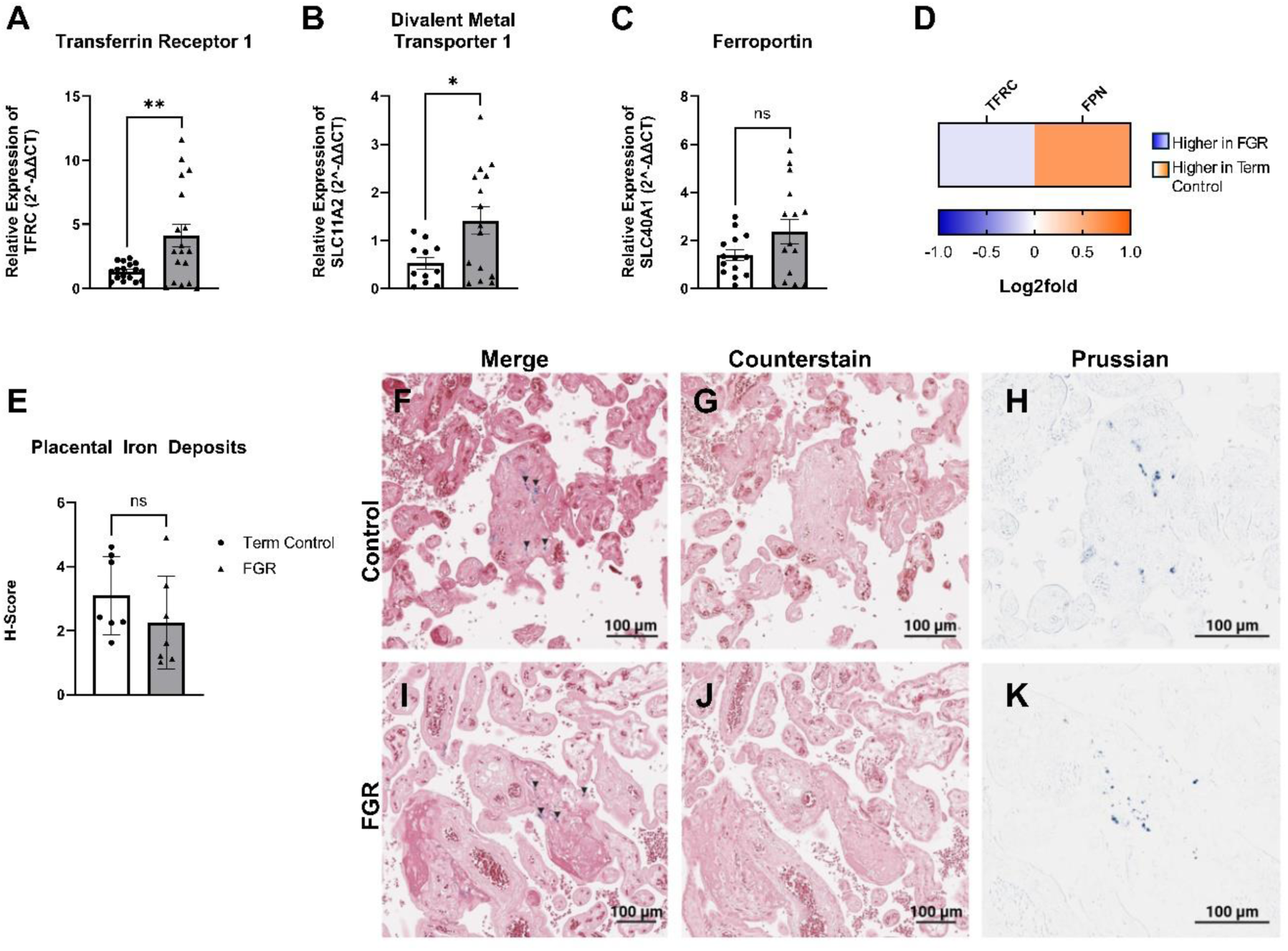
mRNA expression and protein levels of cellular iron transporters and histological Prussian blue staining in FGR and healthy control placental villous tissue. (A) TFRC, Transferrin Receptor 1; (B) SLC11A2, Solute Carrier Family 11 Member 2/ Divalent Metal Transporter 1 (DMT1); (C) SLC40A1, Solute Carrier Family 40 Member 1/Ferroportin (FPN). Data presented as mean ± SEM, control n=11-19 (white and circles), FGR n=10-18 (grey and triangles). Student t-tests were performed, *p<0.05, **p<0.01, ***p<0.001, ****p<0.0001. (D) Proteomes of cellular iron transport depicted as a heatmap based on the log2fold, with transporters indicated on top. The log_2_-FC was derived from the ratio between healthy term placentae to assess changes in FGR placentae, so a positive log_2_-FC indicates higher protein abundance in healthy control placentae (orange), and a negative log_2_-FC indicates higher protein abundance in FGR placentae (blue). (E) Halo Pathology Analysis Software was used to quantify the percentage of Prussian blue staining relative to the tissue area. Heavy, moderate, and light intensity values were used to calculate the H-score in healthy control (n=7) and FGR (n=7) placental tissue. An unpaired t-test revealed no significant difference between the groups. (F) A representative merged image of healthy control placental tissue showing Fe^3+^ deposits (blue staining indicated by arrows). (G) Control placental tissue stained with nuclear fast red counterstain only and (H) Prussian blue staining isolated from counterstained tissue in (F), upon which intensity calculations were performed. (I) A representative merged image of FGR placental tissue, with (J-K) shown similarly to the row above (G-H). The scale bar represents 100μm.

### Mitochondrial Iron Transport

Given the pivotal role of iron in mitochondrial function, we examined mitochondrial iron transporters located within the inner mitochondrial membrane, Mitoferrin 1 (Solute Carrier Family 25 Member 37, SLC25A37) and Mitoferrin 2 (Solute Carrier Family 25 Member 28, SLC25A28). No significant difference in mitoferrin 1 (Figure 2A) mRNA expression was observed in FGR placentae compared to healthy controls. In contrast, mitoferrin 2 (Figure 2B; p<0.005) mRNA expression was significantly lower in FGR placental villous tissue than in healthy controls.

**Figure 2:**
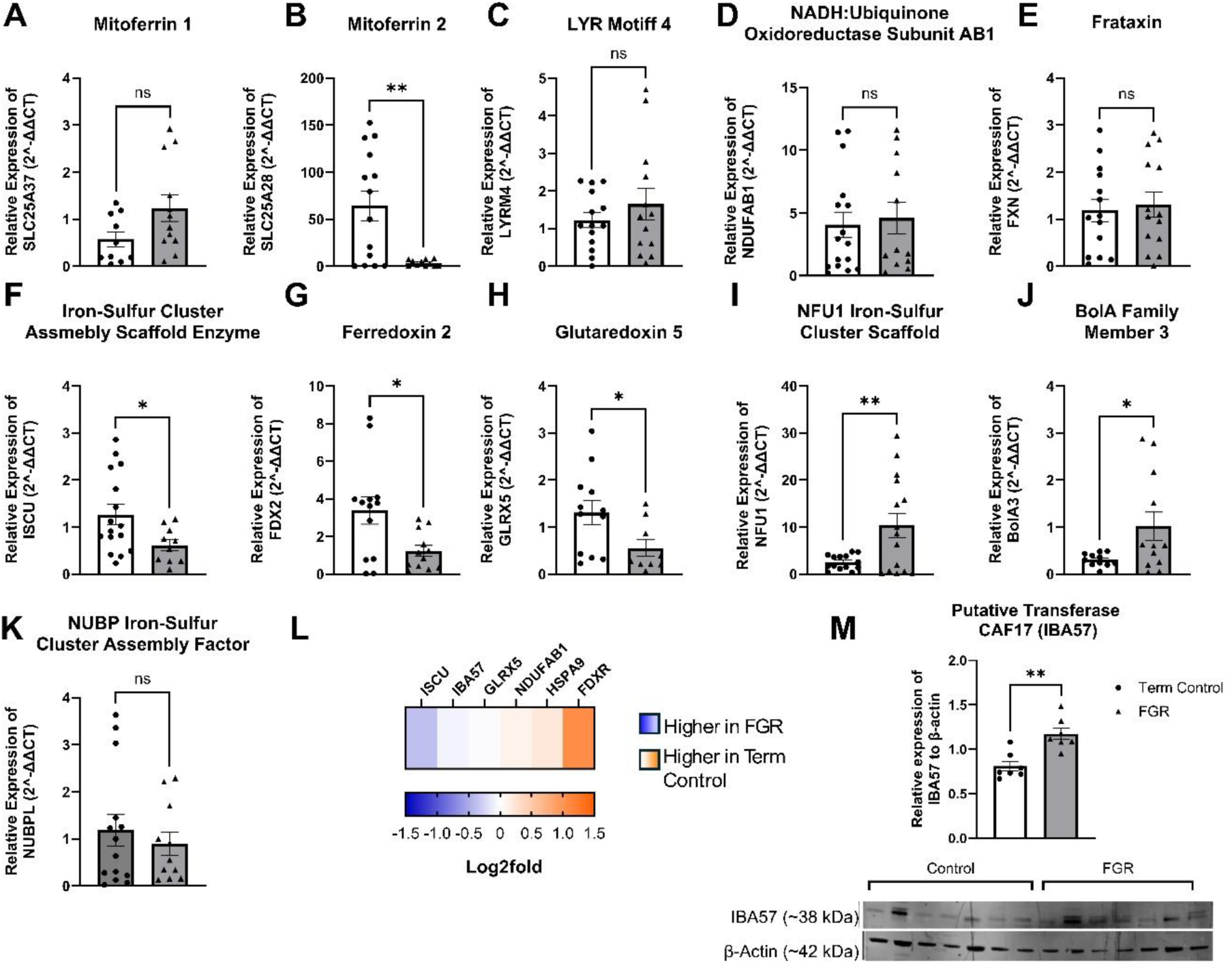
mRNA expression and proteomes of targeted genes in the iron-sulfur (Fe-S) cluster biosynthesis pathway. (A) SLC25A37, Solute Carrier Family 25 Member 37/ Mitoferrin 1; (B) SLC25A28, Solute Carrier Family 25 Member 28/Mitoferrin 2 (C) LYRM4, LYR Motif Containing 4 (ISD11); (D) NDUFAB1, NADH: Ubiquinone Oxidoreductase Subunit AB1; (E) ISCU, Iron-Sulfur Cluster Assembly Enzyme; (F) FXN, Frataxin; (G) FDX1l, Ferredoxin 2 (FDX2); (H) GLRX5, Glutaredoxin 5; (I) NUBPL, NUBP Iron-Sulfur Cluster Assembly Factor; (J) NFU1, NFU1 Iron-Sulfur Cluster Scaffold; (K) BolA3, BolA Family Member 3; Data are presented as mean ± SEM, control n=11-19 (white and circles), FGR n=10-18 (grey and triangles). Student t-tests were performed *p<0.05, **p<0.01, ***p<0.001, ****p<0.0001. (L) Fe-S cluster proteomes, depicted as a heatmap based on log_2_-FC with proteome identification on top. Proteomes of Fe-S cluster assembly depicted as a heatmap based on the log_2_-FC, with transporters indicated on top. The log_2_-FC was derived from the ratio between healthy term placentae to assess changes in FGR placentae, so a positive log_2_-FC indicates higher protein abundance in healthy control placentae (orange), and a negative log_2_-FC indicates higher protein abundance in FGR placentae (blue). (M) Protein abundance of putative transferase CAF17 (IBA57; 38.155kDa) normalised to β-Actin (41.737kDa) in healthy and FGR placentae. Representative immunoblot shown below.

### Mitochondrial Iron Utilisation: De Novo Fe-S Cluster Synthesis

The utilisation of iron within the mitochondria converges on the synthesis of Fe-S clusters. This process comprises two main stages: de novo synthesis of [2Fe-2S] clusters and late-stage genesis of [4Fe-4S] clusters vital to mitochondrial function and cellular viability. In de novo Fe-S cluster synthesis, the mRNA expression of LYR Motif 4 (LYRM4/ISD11; Figure 2C), NADH: Ubiquinone Oxidoreductase Subunit AB1 (NDUFAB1/ACP; Figure 2D) and Frataxin (FXN; Figure 2E) were similar in FGR and healthy control placental villous tissue. Conversely, protein levels of NDUFAB1 (0.123 log_2_-FC; Figure 2L) were lower in FGR placentae compared with healthy controls. Ferredoxin 2 (FDX2/FDX1L) mRNA expression (Figure 2G) was significantly lower (p<0.05), while ferredoxin reductase (FDXR; Figure 2L) was 1.08 log_2_-fold lower in protein levels in FGR compared to controls. The mRNA expression of Iron-Sulfur Cluster Assembly Scaffold Enzyme (ISCU; Figure 2F), which is essential for [2Fe-2S] assembly and transfer, was significantly lower (p<0.05) within FGR placental tissue compared with healthy controls. However, ISCU protein expression (−0.372 log_2_-FC; Figure 2L) was higher in FGR placental villous tissue when compared with healthy controls. An important transporter of the [2Fe-2S] cluster, Glutaredoxin 5 (GLRX5; Figure 2H), had significantly lower mRNA expression (p<0.05) in FGR placentae; however, no difference in protein abundance between FGR and healthy term placentae was observed.

### Late-Stage Fe-S Cluster Synthesis

In late-stage Fe-S cluster assembly, Heat Shock Protein Family A (HSP70) Member 9 (HSPA9; Figure 2L) observed a 0.219 log_2_-fold increase at the protein level in healthy controls compared to FGR placentae. IBA57 (Figure 2L) was −0.073 log_2_-fold higher in protein level with statistical significance (p<0.005) confirmed by western blotting (Figure 2M). [4Fe-4S] cluster transporters showed increased mRNA expression of NFU1 Iron-Sulfur Cluster Scaffold (NFU1; p<0.005, Figure 2I) and BOLA Family member 3 (BOLA3; p<0.05, Figure 2J) in FGR placental villous tissue when compared to healthy controls. No significant changes in mRNA expression were observed in NUBP Iron-Sulfur Cluster Assembly factor (NUBPL; Figure 2K).

### Heme Synthesis

Analysis of the heme synthesis pathway showed significantly higher mRNA expression of coproporphyrinogen oxidase (CPOX; p=0.002; Figure 3A) and ferrochelatase (FECH; p=0.005; Figure 3B) at the gene and protein level, −0.163 log_2_-fold and −0.356 log_2_-fold, respectively (Figure 3D) within FGR placental tissue compared to healthy tissue. Furthermore, protein levels of additional enzymes important to heme synthesis, Uroporphyrinogen Decarboxylase (UROD; Figure 3D) Hydroxymethylbilane Synthase (HMBS; Figure 3D), were −0.175 log_2_-fold and −0.196 log_2_-fold greater in FGR, respectively. In contrast, Aminolevulinate dehydratase (ALAD/PBGS; Figure 3D) was 0.541 log_2_-fold lower protein expression in FGR compared with term control placental villous tissue. Although heme synthesis enzymes were upregulated, this was not conserved when assessing heme activity (p<0.005) in FGR placentae. Heme oxygenase (HMOX) −1 protein abundance was −1.13 log_2_-fold higher in FGR placentae compared to control placentae (Figure 3D). Additionally, NADPH-cytochrome P450 oxidoreductase (POR) protein abundance was −0.157 log_2_-fold higher in FGR placentae than healthy controls.

**Figure 3:**
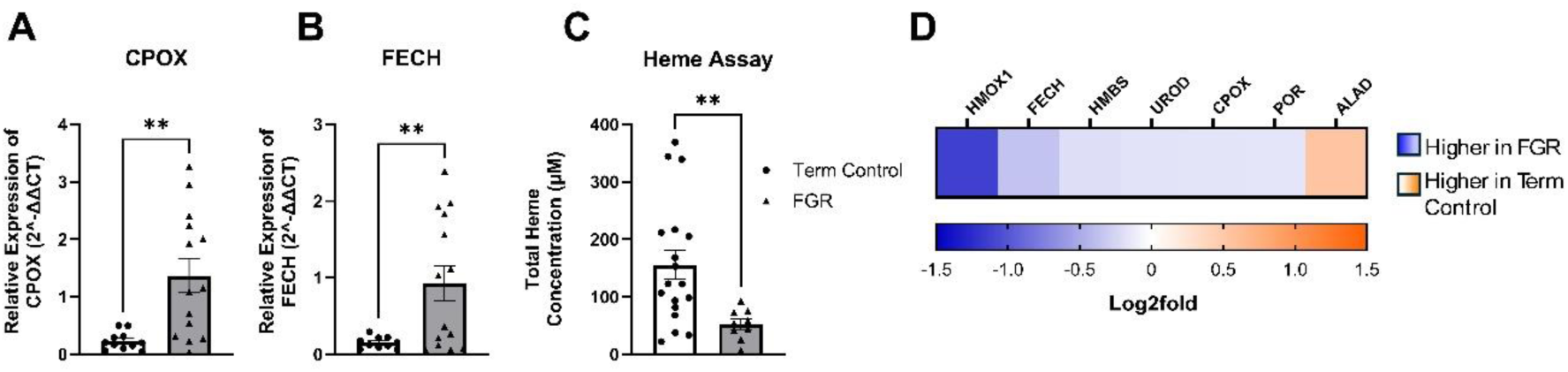
mRNA expression of targeted genes involved in the heme synthesis pathway and heme content in FGR and healthy term placentae. (A) CPOX, Coproporphyrinogen Oxidase; (B) FECH, Ferrochelatase. Data presented as mean ± SEM, control n=11-19 (white and circles), FGR n=10-18 (grey and triangles), with sample loss due to outliers’ tests conducted. Student t-tests were performed *p<0.05, **p<0.01, ***p<0.001, ****p<0.0001. (C) Heme Assay measuring total Heme Concentration (μM) present. (D) Heme synthesis proteomes depicted as a heatmap based on log_2_-FC, with proteome identification on top. Proteomes of heme synthesis depicted as a heatmap based on log_2_-FC, with transporters indicated on top. The log_2_-FC was derived from the ratio between healthy term placentae to assess changes in FGR placentae, so a positive log_2_-FC indicates higher protein abundance in healthy control placentae (orange), and a negative log_2_-FC indicates higher protein abundance in FGR placentae (blue).

The placenta is a potent haemopoietic site where heme synthesis stimulates globin production, forming a haemoglobin tetramer of alpha and beta-like chains critical for binding and transporting oxygen. Our analysis revealed no changes in the mRNA expression of Haemoglobin Subunit Alpha 1 (HBA1) between controls and FGR placental villous tissue, although protein levels were 0.210 log_2_-fold lower in FGR compared to healthy controls (Figure 4A and G). Within the beta-like globin chains, we observed significantly greater mRNA expression of Haemoglobin Subunit Gamma 1 (HBG1; p<0.05, Figure 4B) in FGR placentae, although Haemoglobin Subunit Gamma 2 showed no differences between control and FGR placentae. Protein abundance of HBG1 (0.325 log_2_-FC) and HBG2 (0.092 log_2_-FC) was lower in FGR compared with term control placental tissue samples (Figure 4G). Haemoglobin Subunit Beta (HBB) mRNA expression was significantly higher (p<0.05) in FGR compared to healthy controls (Figure 4D). However, HBB protein abundance was lower (0.585 log_2_-FC) within FGR placental tissue when compared to healthy placental tissue samples (Figure 4G). Levels of Erythrocyte structural proteins, specifically Erythrocyte Protein Band 4.1 (EPB41), EPB42, Spectrin Alpha 1 (SPTA1), Spectrin Beta (SPTB), Ankyrin1 (ANK1) and Solute Carrier Family 4 Member 1 (SLC4A1), were subsequently assessed. The mRNA expression (p<0.05) and protein abundance (0.254 log_2_-FC) of EPB41 were lower in FGR placental tissue compared to healthy tissue controls (Figure 4E and G). EPB42 protein abundance was 0.562 log_2_-fold lower within FGR placentae than healthy placentae (Figure 4G). Analysis of spectrin protein levels, SPTA1 and SPTB (Figure 4G), was 0.156 log_2_-fold and 0.491 log_2_-fold lower in FGR placenta compared with healthy term placenta. Similarly, ANK1 and SLC4A1 protein abundance were 0.661 log_2_-fold and 0.493 log_2_-fold lower in FGR placentae than healthy placentae, suggesting compromised erythrocyte structure. However, analysis of Hemogen (HEMGN; Figure 4F), a regulator of haematopoietic cell differentiation, mRNA expression was similar in healthy and FGR placental villous tissue, indicating that these changes were not occurring at the onset of hematopoiesis but rather at erythrocyte maturation.

**Figure 4:**
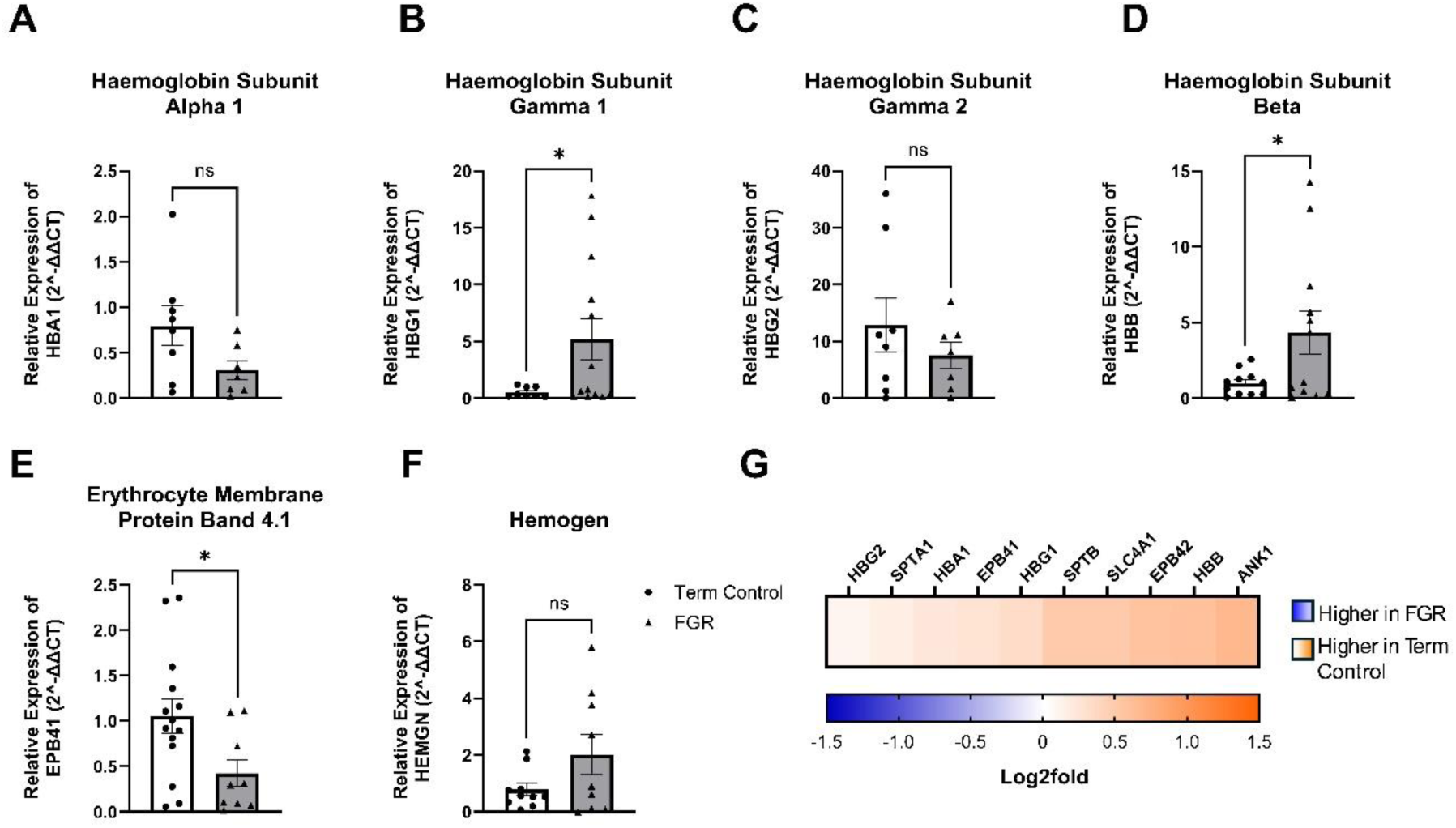
mRNA expression and proteome of targeted genes involved in Haematopoiesis. (A) HBA1. Haemoglobin Subunit Alpha 1; (B) HBG1, Haemoglobin Subunit Gamma 1; (C) HBG2, Haemoglobin Subunit Gamma 2; (D) HBB, Haemoglobin Subunit Beta; (E) EPB41, Erythrocyte Membrane Protein Band 4.1; (F) HEMGN, Hemogen; Data presented as mean ± SEM, control n=11-19 (white and circles), FGR n=10-18 (grey and triangles). Student t-tests test performed *p<0.05, **p<0.01, ***p<0.001, ****p<0.0001.(G) Haematopoiesis proteomes, depicted as a heatmap and based on log_2_-FC with proteome identification on top. Proteomes of haematopoiesis depicted as a heatmap based on log_2_-FC, with transporters indicated on top. The log_2_-FC was derived from the ratio between healthy term placentae to assess changes in FGR placentae, so a positive log_2_-FC indicates higher protein abundance in healthy control placentae (orange), and a negative log_2_-FC indicates higher protein abundance in FGR placentae (blue).

## Discussion

This project sought to elucidate the complex role of iron in placental and mitochondrial function in fetal growth restriction (FGR). As expected, we observed reduced placental and birth weights in FGR compared with healthy controls (Table 2). Similarly, fetal biometric measurements showed significantly reduced abdominal circumference, head circumference, and femur length in FGR pregnancies compared with healthy-term pregnancies. Mothers with FGR babies also showed reduced red cell distribution width (RDW) in conjunction with elevated ferritin levels, although these fell within clinical reference ranges. No changes were observed in circulating maternal iron or haemoglobin levels, and all maternal measurements stayed within the clinical reference ranges (Table 2). Subsequent analysis of FGR placental tissue identified altered iron transporters (Figure 1). Furthermore, FGR placentae showed changes in mitochondrial iron-sulfur (Fe-S) cluster biosynthesis, specifically a reduced capacity for converting [2Fe-2S] to [4Fe-4S] clusters due to lower levels of key [2Fe-2S] proteins ferredoxin reductase (FDXR) and ferredoxin 2 (FDX2) required for synthesis of both [2Fe-2S] and late-stage [4Fe-4S] clusters (Figure 2). The observed reduction in proteins involved in [4Fe-4S] cluster assembly suggests a metabolic redirection from incorporation into the mitochondrial electron transport chain (ETC) toward increased heme production, as evidenced by increased cytosolic and mitochondrial proteins of the heme synthesis proteins (Figure 3). Despite the apparent shift toward heme synthesis, total heme concentrations in FGR placentae were decreased. We propose that this arises from increased regulation of the heme degradation pathway in FGR placentae, evidenced by elevated protein levels of heme oxygenases and NADPH-cytochrome P450 oxidoreductase. Consequently, we observed reduced haemoglobin proteins within the FGR placentae (Figure 4). Additionally, our findings identified alterations in levels of erythrocyte structural proteins in FGR placentae (Figure 4). These alterations may further compromise oxygen delivery to the fetus or simultaneously represent an adaptive response to the poor vasculature of FGR placentae. Together, these findings enhance our understanding of the fundamental biological processes underlying FGR and highlight how adaptations in iron utilisation contribute to mitochondrial dysfunction observed in FGR, perpetuating inadequate fetal growth.

### Maternal Haematological Assessment in FGR

Pregnancy is a dynamic process that is associated with significant alterations in the maternal cardiovascular system to support fetoplacental development (Sanghavi & Rutherford, 2014). Maternal cardiovascular adaptations, including increased plasma volume (20-50%) and red blood cell mass (20-30%), begin rising from week 5 of pregnancy, well before the maternal and fetal circulatory systems are functionally established via the placenta (Allerkamp *et al*., 2022; Mégier *et al*., 2022; Söhnchen *et al*., 2011). Additionally, studies have shown that erythropoiesis is the largest consumer of iron and as gestation advances, the demand for iron increases substantially to support expanded maternal red cell mass, placental development, and fetal growth (Gambling *et al*., 2001; Kautz & Nemeth, 2014). Maternal cardiovascular function and haemodynamic failures have significant implications for the health of both the mother and the fetus.

In FGR pregnancies, normal maternal haemodynamic responses are known to be compromised, including inadequate blood volume expansion and impaired reduction in vascular resistance (Mecacci *et al*., 2021; Perry *et al*., 2020). Therefore, our study assessed whether maternal haemodynamic responses in FGR pregnancies might be reflected in maternal blood parameters (Table 2). We observed reduced maternal red blood cell distribution width (RDW) in FGR mothers compared with controls. RDW is the measurement of variation in red blood cell size and volume relative to the mean red cell volume (MCV) (Fava *et al*., 2019; Shehata *et al*., 1998). Despite being statistically significant, the reduction in maternal RDW remained within clinical reference ranges, indicating decreased anisocytosis, less variability in erythrocyte size and a narrower distribution around the mean cell volume (de Freitas *et al*., 2019; Kurt *et al*., 2015; Lippi & Plebani, 2014). We speculate that the statistical significance in maternal RDW between FGR and healthy control mothers, while within clinical reference ranges, may be a maternal adaptation to overcome poor spiral artery remodelling. Improved erythrocyte uniformity may enhance blood rheology (Nader *et al*., 2019; Reddy *et al*., 2016; Zhang *et al*., 2009), which may provide advantages in the context of increased vascular resistance and narrowed maternal spiral arteries, contributing to the reduced perfusion of the intervillous space often observed in FGR (Burton & Jauniaux, 2018).

Maternal serum ferritin levels were significantly higher in FGR pregnancies than in healthy controls (Table 2), although ferritin levels fell within clinical reference ranges (30-200 μg/L). Our observations are substantiated by previous studies showing that elevated serum ferritin levels increased the risk of developing FGR (Ozgu-Erdinc *et al*., 2014). Furthermore, Hou *et al*. (2000) demonstrated that FGR pregnancies with elevated serum ferritin levels presented with asymmetric fetal growth during the second and third trimesters. Asymmetrical growth is a key hallmark of late-onset FGR (diagnosis >32 weeks), accounting for 70-80% of FGR cases (Sun *et al*., 2020). This may explain why serum ferritin levels were higher in FGR cases within our study, as 83% of our samples were delivered >32 weeks. Moreover, serum ferritin’s specificity (true negative rate in identifying iron sufficiency) and sensitivity (true iron deficiency) are essential for assessing biological iron status when chronic inflammation is absent (Mégier *et al*., 2022). The increase in maternal serum ferritin may, therefore, be attributed to an inflammatory response as ferritin acts as an acute-phase reactant during inflammation (Camaschella, 2015), rather than truly indicating maternal iron status, as pregnancy is inherently pro-inflammatory. This is supported by increases in oxidative stress and inflammation observed in the maternal circulation and placenta of FGR pregnancies (Hou *et al*., 2000; Ozgu-Erdinc *et al*., 2014).

Despite statistically elevated maternal serum ferritin levels and reduced RDW in FGR, the lack of maternal changes in iron haematological parameters, outside of normal reference ranges, suggests that subsequent findings were not due to impaired iron delivery but rather demonstrate that maternal adaptations are likely secondary responses to compromised placental function.

### Iron transport in the FGR placenta

Our study revealed that iron transporter proteins were altered in FGR placentae, characterised by an enhanced capacity for iron uptake. Specifically, we observed a 1.08-fold increase in Transferrin receptor 1 (TfR1/TFRC) levels (−0.117 log2-fold), the major importer of non-heme iron into cells, in FGR placenta (Figure 1A and D). While maternal iron deficiency has been shown to increase TfR1 expression in placental tissue, ensuring appropriate iron uptake from maternal circulation to support fetal and placental iron requirements, we found no evidence of maternal iron deficiency within our FGR group (Table 2). However, Burton and Fowden (2012) and Sibley *et al*. (2010) report that when placental size and capacity are reduced, the placenta increases nutrient transporter activity as a compensatory mechanism to suboptimal conditions, rather than failing. This may explain our observed increases in iron transporters in significantly smaller FGR placentae (Table 2). Collectively, this suggests that the increase in TfR1 in our FGR placentae might represent a compensatory response to maximise iron uptake in a compromised placental vascular environment.

Since iron transport is transcellular and unidirectional from mother to fetus, the placenta serves as the “gatekeeper” for iron transfer to the fetus (O’Brien, 2022; Roberts *et al*., 2020). The fetus depends entirely on maternal iron transfer through the placenta, mediated by ferroportin (FPN), which exports iron into the fetal circulation (Figure 1C-D) (Sangkhae *et al*., 2020). We observed 1.56-fold lower FPN (0.646 log2-fold) levels in FGR placental tissue, suggesting that the placenta is retaining iron for its own use (O’Brien, 2022; Sangkhae *et al*., 2020). This is supported by an increase in divalent metal transporter 1 (DMT1/SLC11A2) mRNA expression, the transporter of iron into the cell cytoplasm following endocytosis of the TfR1-iron complex (Figure 1B). Subsequent Prussian blue staining showed no difference in iron deposit content for ferric (Fe^3+^) iron between FGR and healthy controls. Collectively, these findings suggest that iron is being prioritised for intracellular function in FGR placentae, rather than for storage or being exported into fetal circulation.

### Mitochondrial Iron Transport

The mitochondria utilise between ∼20-50% of cellular iron (Qi *et al*., 2023), facilitating integral mitochondrial function and bioenergetic pathways to support fetoplacental growth (Aye *et al*., 2022; Paul *et al*., 2017). Mitoferrin-1 (SLC25A39) and mitoferrin-2 (SLC25A28) facilitate iron transport across the inner mitochondrial membrane but exhibit different cell expression profiles and cell-specific functions (Figure 2A-B) (Ali *et al*., 2022). We observed no changes in mitoferrin-1 mRNA expression between our groups. In line with published literature in mice and zebrafish haematopoietic tissue (liver, bone marrow, spleen, and yolk sac islands) and embryonic tissue that determined mitoferrin-1 is essential for erythroid development to meet the higher demand for mitochondrial iron to support heme production (Paradkar *et al*., 2009; Shaw *et al*., 2006). As the placenta is a potent haematopoietic tissue containing erythroid progenitors from as early as 6 weeks of gestation through to term (Gekas *et al*., 2005; Robin *et al*., 2009). We propose that the expression of mitoferrin-1 is likely conserved, attributed to the placenta’s fundamental role as a haematopoietic organ, rather than solely as a facilitator of maternal-fetal exchange. This is further demonstrated by Shaw *et al*. (2006) who observed that knockout (KO) of mitoferrin-1 in murine models results in embryonic lethality at the initiation of definitive erythrocyte development.

In contrast to mitoferrin-1, which is predominantly expressed in erythroid tissue, and minimal expressed in other tissues (Ali *et al*., 2022), Shaw *et al*. (2006) and Paradkar *et al*. (2009) found that mitoferrin-2 is ubiquitously expressed in non-erythroid tissue and is responsible for the iron import required for haemoproteins and Fe-S cluster synthesis (Ali *et al*., 2022). Our study observed a decrease in mitoferrin-2 mRNA expression in placentae from FGR pregnancies. These findings highlight the complex regulation of iron transport in FGR placentae, suggesting a selective downregulation of mitoferrin-2 while preserving mitoferrin-1 iron uptake for haematopoietic use.

### Altered Iron-Sulfur Cluster Biosynthesis Machinery in FGR Placental Mitochondria

Given that our data suggested a conservation of intracellular iron and altered mitochondrial iron uptake within FGR placentae, we went on to investigate iron-sulfur (Fe-S) cluster biosynthesis. Fe-S clusters are apoproteins essential for cellular activities, including heme and are directly incorporated into the mitochondrial electron transport chain (ETC) to support mitochondrial structure and function (Guan *et al*., 2022; Voltarelli *et al*., 2023). Fe-S assembly is a complex process that can be categorised into two main components: (i) de novo synthesis of [2Fe-2S] cluster and (ii) late Fe-S ([4Fe-4S]) cluster synthesis (Figure 3). Disruptions in Fe-S biogenesis impair cellular and mitochondrial function, and have been shown in literature to lead to metabolic myopathies such as Friedreich’s ataxia, sideroblastic anaemia, and ISCU myopathy (Lee *et al*., 2011).

### Mitochondrial De Novo Fe-S Synthesis in FGR

De novo Fe-S cluster synthesis is a complex series of reactions, initiated by the catalysis of a pyridoxal-phosphate, cysteine desulfurase NFS1 (Maio & Rouault, 2020). Stabilising the activity of NFS1, NADH: Ubiquinone Oxidoreductase Subunit AB1 (NDUFAB1) forms a heterodimer with LYR Motif Containing 4 (LYRM4), initiating Fe-S cluster synthesis via the complex NFS1:LYRM4:NDUFBA1 (Figure 2C-D) (Herrera *et al*., 2019). Our data showed a 1.09-fold lower abundance of NDUFAB1 (0.123 log_2_-fold) in FGR placentae, despite the heterodimer conjugate LYRM4 mRNA expression remaining unchanged. The subsequent synthesis of the NFS1:LYRM4:NDUFBA1 complex results in the binding of the iron-sulfur cluster assembly enzyme (ISCU) and frataxin (FXN), forming a hetero-octamer that consists of two copies of the NFS1:LYRM4:NDUFBA1:FXN:ISCU complex (Figure 2C-F and L) (Maio & Rouault, 2020).

In our study, FXN mRNA expression remained unchanged between the two groups (Figure 2E), suggesting any deficits in Fe-S assembly likely occur downstream of initial FXN-mediated iron and sulfide delivery onto ISCU, as loss of FXN results in widespread depletion of Fe-containing proteins and attenuation of mitochondrial protein synthesis (Figure 2E) (Zhong *et al*., 2023). We observed a decrease in ISCU mRNA, which provides the structural scaffold for [2Fe-2S] formation, in FGR placental mitochondria (Figure 2F). However, this observation was not conserved in our proteomics data, with a 1.29-fold higher protein abundance in FGR (−0.372 log_2_-FC; Figure 2L). In addition, key [2Fe-2S] proteins ferredoxin reductase (FDXR) and ferredoxin 2 (FDX2), which are required to supply electrons to complete [2Fe-2S] and [4Fe-4S] assembly, had 2.11-fold lower protein (1.08 log_2_-FC) and decreased gene expression (p<0.05) in FGR placentae, respectively. As we did not observe global suppression, we suggest the FGR placenta prioritises [2Fe-2S] biogenesis at the detriment of [4Fe-4S] cluster synthesis (Figure 2G and L). This is consistent with previous knockdown studies showing that loss of FDX2 globally disrupts mitochondrial Fe-S assembly, leading to dysregulated oxidative phosphorylation (OXPHOS) and reduced antioxidant production (Shi *et al*., 2012).

Following [2Fe-2S] cluster formation, GLRX5 receives the [2Fe-2S] cluster from ISCU for transport to apoproteins required for various cellular and mitochondrial functions (Paul *et al*., 2017). We observed a significant decrease in mRNA expression of GLRX5, yet GLRX5 protein abundance remained unchanged between the two groups (Figure 2H and L), again preserving the capacity for [2Fe-2S] cluster transfer to downstream cellular and mitochondrial functions such as heme synthesis or late-stage Fe-S cluster formation.

### Mitochondrial Late-Stage Fe-S Synthesis in FGR

Given the changes observed in de novo synthesis, specifically reductions in protein abundance of FDXR and mRNA expression of FDX2 and their sequential roles in electron transfer in the formation of [2Fe-2S] and [4Fe-4S] clusters, we investigated late-stage Fe-S synthesis. In contrast to de novo Fe-S cluster synthesis, late-stage [4Fe-4S] clusters are critical cofactors that are incorporated into mitochondrial ETC complexes, and apoproteins such as iron-regulatory proteins, and antioxidants (Maio & Rouault, 2020).

The formation of [4Fe-4S] clusters requires the transfer of [2Fe-2S] intermediaries to the A-type scaffold proteins, Iron-Sulfur Cluster Assembly 1 and 2 (ISCA1 and ISCA2) in conjugation with Iron-Sulfur Cluster Assembly Factor IBA57 (Melber *et al*., 2016; Paul *et al*., 2017). In FGR placentae, we observed a small 1.05-fold increase in IBA57 protein (−0.073 log_2_-FC; Figure 2L), which was confirmed by western blotting to be significant (p<0.002; Figure 2M). However, we did not detect ISCA1 and ISCA2 at sufficient levels within our proteomic analysis. Given that ISCA1 and ISCA2 are known to contain [2Fe-2S] clusters (Beilschmidt *et al*., 2017; Nasta *et al*., 2019) but IBA57 does not contain [2Fe-2S] clusters (Amick *et al*., 2014; Dores-Silva *et al*., 2015), the inability to detect sufficient levels of ISCA proteins in FGR may suggest that the newly synthesised [2Fe-2S] clusters cannot be effectively matured into [4Fe-4S] clusters.

Newly synthesised [4Fe-4S] clusters are released from the ISCA1:ISCA2:IBA57 complex by a chaperone/co-chaperone system comprising heat shock protein family A (HSP 70) member 9 (HSPA9) and mitochondrial Fe-S Cluster co-chaperone (HSC20). HSPA9 protein abundance was 1.16-fold lower in FGR placentae (0.219 log_2_-FC; Figure 2L). Following [4Fe-4S] cluster release, HSPA9 mediates the transfer [4Fe-4S] clusters directly to apoproteins or through targeting factors such as NFU1 Iron-Sulfur Cluster Scaffold (NFU1), BolA Family Member 3 (BOLA3) and NUBP Iron-Sulfur Cluster Assembly Factor (NUBPL) (Maio & Rouault, 2020; Melber *et al*., 2016; Zhong *et al*., 2023). Incorporation of [4Fe-4S] into the mitochondrial ETC complexes provides electron tunnelling through complexes I-IV, generating energy and a proton gradient crucial for ATP synthesis (Mostajabi Sarhangi & Matyushov, 2023; Vanlander & Van Coster, 2018). Despite increased mRNA expression of BOLA3 and NFU1 in FGR placentae (Figure 2I-J), we observed no changes in NUBPL mRNA expression (Figure 2K), which may suggest an attempt to maximise the delivery of a limited pool of [4Fe-4S] clusters.

Collectively, the reduced FDX2 mRNA expression and lower abundance of FDXR and HSPA9 suggest a bottleneck in mitochondrial Fe-S cluster biogenesis in the FGR placenta. While these proteins participate in both [2Fe-2S] and [4Fe-4S] cluster formation (Maio & Rouault, 2020), their decreased abundance in FGR likely prioritises the direction of limited resources toward essential [2Fe-2S] cluster formation, for heme synthesis (Shi *et al*., 2012; Weiler *et al*., 2020). This disruption in mitochondrial Fe-S assembly provides a molecular basis for the mitochondrial dysfunction observed in FGR placentae, characterised by impaired respiratory chain complex activity, altered ATP production, and increased oxidative stress (Holland *et al*., 2017; Xu *et al*., 2021).

### Altered Heme Synthesis and Erythrocyte Structure in FGR Placentae: Implications for Fetal Oxygenation and Development

Based on our data showing prioritisation of [2Fe-2S] clusters in FGR placentae, we further investigated the heme synthesis pathway. Heme synthesis critically depends on the insertion of [2Fe-2S] clusters into ferrochelatase (FECH), the terminal enzyme of heme biosynthesis that resides within the mitochondria. The insertion of [2Fe-2S] clusters into FECH ensures its stability and activity for heme production (Al-Karadaghi *et al*., 1997; Obi *et al*., 2022). Our data revealed significantly higher levels of enzymes across the heme synthesis pathway in FGR placentae (Figure 3D). Specifically, elevated mRNA and protein levels of enzymes coproporphyrinogen oxidase (CPOX; 1.12-fold; −0.163 log_2_ FC) and FECH (1.28-fold; −0.356 log_2_-fold), located within the mitochondria, were accompanied by greater protein abundance of heme enzymes Hydroxymethylbilane Synthase (HMBS; 1.15-fold), and Uroporphyrinogen Decarboxylase (UROD: 1.13-fold). Supporting our suggestion that the FGR placentae prioritises [2Fe-2S] clusters and shifts utilisation towards heme synthesis. Additionally, FECH has been reported to be complexed to ATP Binding Cassette Subfamily B Member 10 (ABCB10), conferring stability to mitoferrin-1. This FECH-ABCB10 conjugation may explain why mitoferrin-1 expression remains conserved in FGR placentae despite reduced mitoferrin-2.

Beyond its role in oxygen transport and haemoglobin production, heme also contributes to mitochondrial respiration by facilitating electron transfer and proton translocation within ETC complexes II, III, and IV through its incorporation as heme a, b, and c moieties (Kim et al., 2012). Despite the increased levels of heme synthesis enzymes, the assessment of heme concentration in FGR placentae was lower (Figure 3C). This may be due to increased incorporation of heme a, b and c into ETC complexes II, III and IV to increase mitochondrial respiration resulting from a lack of [4Fe-4S] assembly, although this hypothesis requires confirmation.

Alternatively, heme is a known pro-oxidant, inducing oxidative stress, and heme accumulation can induce its catabolism via heme oxygenase 1 (HMOX1), degrading heme into carbon monoxide, iron and biliverdin (Figure 3D). We observed 2.19-fold higher HMOX1 abundance in FGR placentae, suggesting that despite upregulation of the heme biosynthesis pathway, increased heme breakdown may reflect heightened oxidative stress or a compensatory mechanism to regulate “free” heme levels. The catabolism of heme by the placenta is not well understood (Liu et al., 2016), although Peoc’h et al. (2020) have suggested that HMOX expression in the placenta could be a protective response against oxidative stress, which may explain why HMOX1 was higher in FGR placentae.

Our findings suggest a delicate balance in placental heme metabolism in FGR, whereby the upregulation of heme synthesis pathways likely represents an adaptive response to fetoplacental hypoxia and maintenance of placental bioenergetics. Simultaneously, the increase in HMOX1-mediated heme catabolism serves as a protective mechanism against heme-induced oxidative stress.

### Haematopoietic function in FGR placentae

In addition to its role in mitochondrial function, heme is integral for the production of various globin chains, including gamma (γ), alpha (α) and beta (β) (Figure 4). As we observed an increase in heme-producing proteins, but no corresponding change in available heme in our FGR placental tissue, we examined the intricate relationship between heme and globin synthesis. This fundamental process is central to the oxygen-carrying capacity of blood and, by extension, aerobic respiration essential for cellular function and viability (Hamza & Dailey, 2012; Voltarelli et al., 2023). During embryonic development, primitive embryonic erythrocytes are located in blood islands of the secondary yolk sac (Manning *et al*., 2020). Subsequently, a crucial transition occurs, coinciding with placental formation and the emergence of definitive erythropoiesis (Palis & Segel, 1998). This developmental shift results in the production of fetal haemoglobin (α2γ2), which becomes the predominant form throughout the remainder of fetal development (Hardison, 2012). This haemoglobin switching process represents a critical milestone in fetal haematological development and is tightly regulated by complex genetic and epigenetic factors (Hardison, 2012; Manning *et al*., 2020). Fetal haemoglobin production begins at around 10 weeks of gestation, peaking at 20 weeks and remaining constant until the final month of fetal development when gamma-globin synthesis rapidly decreases (Cantú & Philipsen, 2014; Mavilio *et al*., 1983). Our findings revealed lower protein abundance of gamma-1 (HBG1; 1.25-fold; 0.325 log_2_-FC) and gamma-2 (HBG2: 1.09-fold; 0.09 log_2_-FC), key beta-like globin components of fetal haemoglobin, along with haemoglobin subunit alpha 1 (HBA1; 1.16-fold; 0.21 log_2_-FC), in FGR placental tissue compared to healthy term placental tissue (Figure 4). Additionally, the mRNA expression of haemoglobin subunit beta (HBB; p<0.05), a critical component of adult haemoglobin (α_2_β_2_), was significantly increased in FGR placentae, however, protein abundance for HBB was 1.50-fold lower (Figure 4D and G). The presence of HBB in the placenta aligns with current literature stating that β-globin synthesis begins during the last month of fetal development (Cantú & Philipsen, 2014). The switch from fetal to adult haemoglobin occurs when erythropoiesis shifts from the fetal liver to the bone marrow at approximately (Khandros & Blobel, 2024).

While fetal hypoxia in FGR is typically associated with elevated erythropoietin (EPO) production and increased fetal haemoglobin synthesis (Chang *et al*., 2018), our findings indicate a disruption in globin chain production. The lower protein abundance of key globin subunits, HBG1, HBG2, HBB and HBA1, in FGR placental tissue, suggests that globin chain synthesis or stability may be impaired, potentially limiting the assembly of functional haemoglobin. Although recent studies describe the placenta as a transient hematopoietic niche, particularly during early to mid-gestation (Ivanovs *et al*., 2017; Rhodes *et al*., 2008), this function may be compromised in FGR. Chronic hypoxia, inflammation, and oxidative stress, all hallmarks of FGR pathophysiology, can disrupt haematopoietic stem and progenitor cell (HSPC) function and erythroid differentiation (Dzierzak & Bigas, 2018). While hypoxia is known to drive the upregulation of fetal haemoglobin via HIF1α activation (Khandros & Blobel, 2024), we instead observed lower levels of subunits in both fetal and adult haemoglobin. Together, suggesting a dysfunctional placental erythropoietic environment in FGR.

Given the substantial alterations in both heme and globin subunits, we further investigated the erythropoietic structure in the FGR placenta. Our study revealed significant alterations in key structural proteins within FGR placentae. Notably, we observed reductions in erythrocyte membrane protein band 4.1 (EPB41) expression at both mRNA (p<0.05) and protein levels (1.19-fold; 0.253 log_2_-FC) in FGR placentae (Figure 4E and G). The crucial role of EPB41 is in stabilising the actin-spectrin cytoskeleton of erythrocytes, thereby maintaining the characteristic biconcave disc shape of erythrocytes. The biconcave shape is fundamental to erythrocytes’ function, optimising surface area to volume ratio and facilitating efficient oxygen diffusion and laminar flow in circulation (Uzoigwe, 2006; Vorn *et al*., 2023). Huang et al (2017) showed that EPB41 null mice failed to develop a biconcave shape, resulting in compromised oxygen diffusion capacity. Furthermore, we observed reduced protein abundance in EPB42 (1.48-fold; 0.562 log_2_-FC), SPTA1 (1.12-fold; 0.153 log_2_-FC), SPTB (1.40-fold; 0.0491 log_2_-FC), ANK1 (1.57-fold; 0.661 log_2_-FC) and SLC4A1 (1.40-fold; 0.493 log_2_-FC) in FGR placentae when compared to healthy controls (Figure 4G). Similarly, these proteins are integral components of the erythrocyte membrane, conferring flexibility and stability (Alper, 2009; Guo *et al*., 2022; Vorn *et al*., 2023) Mutations in the aforementioned genes have been implicated in hereditary spherocytosis (Xu *et al*., 2023), a condition characterised by erythrocyte membrane instability and deformation. Our findings suggest altered erythropoietic structure during erythrocyte maturation, as HEMGN mRNA expression remained unchanged, indicating that the HSPC population and early erythroid commitment remain preserved.

We propose that changes in the erythropoietic structure reflects a hypoxia-driven alteration in erythroid maturation, skewing erythropoiesis toward a rapid, stress-induced phenotype regulated in part by HIFs (Haase, 2013). Hypoxia-induced oxidative stress, as characteristic of FGR (Xu *et al*., 2021), has been shown to affect the post-translational folding or stability of erythrocyte membrane components, which may further affect the abundance of EPB41 and EPB42 (Mohandas & Gallagher, 2008) and other erythrocyte membrane-bound proteins. Stress-induced erythropoiesis favours accelerated erythropoiesis, which may compromise erythrocyte cytoskeletal assembly and membrane protein integration (Paulson *et al*., 2020). Collectively, these alterations may represent an attempt to increase erythrocyte deformability to facilitate circulation within the structurally compromised vasculature of the FGR placenta, characterised by narrower blood vessels and increased vascular resistance (Boss *et al*., 2023; Kovo *et al*., 2013; Sun *et al*., 2020). Thus, enabling efficient uptake and transport of the limited oxygen available to the fetus.

Despite the structural and functional changes observed in erythrocytes of FGR placental tissue, the mRNA expression of hemogen (HEMGN) remained unchanged. Hemogen regulates the proliferation and differentiation of haematopoietic cells (Jiang *et al*., 2023) indicating that the architectural insult observed in FGR placentae is likely not occurring during haematopoiesis but rather during erythrocyte maturation (Vorn *et al*., 2023) or as a result of stress-induced erythrocyte maturation (Paulson *et al*., 2020).

## Limitations

While previous research indicates that maternal iron status is a critical determinant of fetal iron acquisition through placental transport (Cao & Fleming, 2016; Fisher & Nemeth, 2017; Roberts *et al*., 2020), our study has provided new insights into intrinsic placental changes that are independent of maternal iron status, revealing potential intracellular alterations that have remained elusive in the development of FGR. Since, the placenta is the critical interface between maternal and fetal circulations, building upon the biomolecular foundation established in this study incorporating fetal cord blood analysis in future studies would provide further insight into how modifications in placental iron transport and utilisation impact fetal iron status. In this study, maternal patient records were retrieved from mothers who delivered either healthy or FGR infants, offering valuable insights into maternal iron status during pregnancy. The clinical records revealed comparable iron parameters between groups, strengthening our conclusion that the placental iron alterations we observed represent intrinsic placental adaptations rather than responses to maternal iron availability. Although the retrospective nature of clinical records means that iron parameters were measured based on clinical need rather than a standardised gestational timepoint. While this approach provides real-time data collected during routine clinical care, data uniformity across participants may be potentially limiting (Talari & Goyal, 2020).

This study employed a bottom-up liquid chromatography mass spectrometry (LC-MS) proteomic approach to successfully identify significant alterations in placental iron transport and mitochondrial iron utilisation. While the LC-MS methodology chosen is well-suited to global profiling performed within this study to assess total cellular protein content, some proteins were difficult to detect due to protein inference ambiguity, underrepresentation of low-abundance, membrane-bound proteins, and incomplete coverage of post-translational modifications (Bruderer *et al*., 2015; Mulhall *et al*.). Notably, key Fe-S cluster proteins essential for de novo and late-stage synthesis were difficult to detect, likely due to their hydrophobic nature and the inherent limitations of the protocol (Mulhall *et al*.). Future studies incorporating targeted approaches may improve the detection and quantification of such proteins.

Our study primarily focused on non-heme iron pathways within the placenta, which represents 85% of iron obtained from the maternal diet through plant-based sources of iron (Piskin *et al*., 2022; Yiannikourides & Latunde-Dada, 2019). Heme-iron, accounting for 15% of dietary iron from animal-based sources, requires further investigation to understand heme-iron absorption during pregnancy. Current literature suggests that heme-iron absorption follows the same pattern as non-heme iron absorption in the placenta and is mainly sourced via maternal haemolysis rather than absorbed dietary iron (Milman, 2006; Roberts *et al*., 2020).

## Conclusion

Our study reveals specific alterations in placental iron transport and utilisation pathways in FGR, that are independent of maternal iron status. We identified increased placental iron transporters alongside reduced ferroportin, suggesting an adaptive iron retention within FGR placentae. Assessment of mitochondrial iron utilisation pathways in FGR placentae suggested a preference for [2Fe-2S] cluster formation and heme synthesis over [4Fe-4S] cluster assembly in FGR. Collectively, these findings of altered placental iron transporters and modifications in mitochondrial Fe-S cluster and heme synthesis pathways likely reflect the placenta’s response to the compromised vascular environment characteristic of FGR. We also observed reduced haemoglobin subunits and changes in erythrocyte structural proteins in FGR placentae, indicative of altered placental hematopoietic function. We propose these data are a programmed response to hypoxia and stress-induced haematopoiesis to overcome the poor placental vasculature in FGR. Together, these findings provide new insights into how alterations in iron transport and iron-dependent pathways contribute to the development of FGR.

## Data Availability Statement

The authors will make the raw data supporting the conclusions of this article available, without undue reservation, to any qualified researcher.

## Author Contributions

VBB, KGP, RS and JJF contributed to the conception and design of the study. VBB, HM and SA contributed to the acquisition, analysis and interpretation of data for the study. VBB, KGP, RS, and JJF contributed to drafting the work and critically revising the intellectual content. All authors have approved the final version of the manuscript. All authors agree to be accountable for all aspects of the work. All persons designated as authors qualify for authorship and are listed.

## Acknowledgements

We gratefully acknowledge the research midwives at John Hunter Hospital, Skye Doel and Bridget McCleery, who were involved in participant recruitment and clinical data collection, for their invaluable support and contribution to this study.

